# Schizophrenia-related cognitive dysfunction in the Cyclin-D2 knockout mouse model of ventral hippocampal hyperactivity

**DOI:** 10.1101/301226

**Authors:** Christina M. Grimm, Sonat Aksamaz, Stefanie Schulz, Jasper Teutsch, Piotr Sicinski, Birgit Liss, Dennis Kätzel

## Abstract

Elevated activity at the output stage of the anterior hippocampus has been described as a physiological endophenotype of schizophrenia, and its development maps onto the transition from its prodromal to its psychotic state. Interventions that halt the spreading glutamatergic over-activity in this region and thereby the development of overt schizophrenia could be promising therapies. However, animal models with high construct validity to support such pre-clinical development are scarce. The Cyclin-D2 knockout (CD2-KO) mouse model shows a hippocampal Parvalbumin-interneuron dysfunction and its pattern of hippocampal over-activity shares similarities with that seen in prodromal patients. In a comprehensive phenotyping of CD2-KO mice, we found that they displayed novelty-induced hyperlocomotion (a correlate in the positive symptom domain), that was largely resistant against D1- and D2-dopamine receptor antagonism, but responsive to the mGluR2/3-agonist LY379268. In the negative symptom domain, CD2-KO mice showed transiently reduced sucrose-preference (anhedonia), but enhanced interaction with novel mice and objects, as well as normal nest building and incentive motivation. Also, unconditioned anxiety, perseveration, and motor impulsivity were unaltered. However, in the cognitive domain, CD2-knockouts showed reduced executive function in assays of rule-shift and rule-reversal learning, but also an impairment in working memory, that was resistant against LY379268. In contrast, sustained attention and forms of spatial and object-related memory that are mediated by short-term habituation of stimulus-specific attention were intact. Our results suggest, that CD2-KO mice are a valuable model in translational research targeted at the pharmacoresistant cognitive symptom domain in causal relation to hippocampal over-activity in the prodrome-to-psychosis transition.

## INTRODUCTION

The specific re-creation of endophenotypes of psychiatric diseases in animal models has been proposed as a promising strategy to meet the current challenge of identifying rodent models with predictive validity for drug discovery ^1^. In schizophrenia, hyperactivity of the output stages of the anterior hippocampus represents a physiological endophenotype that is unique in predicting disease progression from the prodromal state to overt schizophrenia (psychosis) ^2,3^: baseline metabolic hyperactivity in the anterior *CA1* region is elevated in prodromal patients that will develop psychosis in the near future, compared to individuals that do not. The development of hyperactivity in the anterior *subiculum*, the output stage of the hippocampus that is excited by CA1, in turn, coincides with the onset of psychosis and is accompanied by atrophy of the anterior hippocampus, which might manifest the disease state irreversibly ^2^. Therefore, interventions that ameliorate anterior hippocampal hyperactivity in the prodromal state and thereby halt further disease progression could be a promising therapeutic strategy ^4^

To support this research, rodent models that display hyperactivity of the ventral hippocampus (vHC, which is homologous to the human anterior hippocampus) are necessary. In rats, rodent correlates of the positive symptom domain were induced by direct electrical or chemical stimulation of the vHC ^5–10^, but also by developmental manipulations such as pre-natal methylazoxymethanol acetate (MAM) injections ^11–13^, which lead to vHC hyperactivity later (but also prefrontal aberrations). It is well established, that such vHC hyperactivity entails a hyper-dopaminergic state of the mesolimbic projection and thereby deficits of latent and pre-pulse inhibition and enhanced novelty- and amphetamine-induced hyperlocomotion.

Other studies focused largely on the contribution of a hypofunction of inhibitory interneurons to vHC hyperactivity. It had been suggested previously, that a homeostatic downregulation of the GABAergic output from Parvalbumin-positive (PV) interneurons of the CA1-region of the hippocampus might lead to an effective disinhibition of the pyramidal neurons of CA1, which would then over-activate the excitatory projection neurons of the ventral subiculum, that project to multiple target areas in the brain that are relevant for affective and cognitive functions ^14^ Chemogenetic silencing of vHC interneurons in mice ^15^ as well as pharmacological vHC disinhibition with picrotoxin in rats ^16^ could indeed provoke some impairments in the positive and the cognitive symptom domains (especially hyperlocomotion and deficits of spatial short-term memory).

A different set of studies focused on the Cyclin-D2 knockout mouse model *(Ccnd2^−/−^*, or CD2-KO)^17^ that displays a reduced number of PV-cells in the hippocampus (20–40 % reduction), particularly in CA1, and some areas of the neocortex (10–30 °%), but not the medial prefrontal cortex ^18,19^. While the deletion of *Ccnd2* affects cell development in the subventricular zone of the medial ganglionic eminence where PV-cells originate from, Somatostatin-positive interneurons that derive from the same region, are not decreased in their numbers in the adult CD2-KO brain ^19^. Physiologically, these animals show hyperactivity of the dentate gyrus, CA2/CA3 and CA1 of the vHC ^18^ – but not the ventral subiculum; a pattern that is more akin to the prodromal state (see above) in which patients already present with cognitive symptoms. However, schizophrenia-related cognitive deficits have not been systematically examined in this mouse line yet (see Table 1 for a summary on previous behavioural assessments of *Ccnd2^−/−^* mice) neither in most other rodent models of ventral hippocampal hyperactivity ^16,20^. However, CD2-KO mice display less efficiency in their search strategies in the water-maze, and a deficit in spatial forms of cognitive flexibility, while basic spatial learning is intact ^21–24^

**Table 1:** Overview over previous behavioural studies in CD2-KO mice. FC, fear-conditioning. Numbers refer to citations in the reference list.

| **Psychological Function** | **Behavioural test** | **Phenotype** |
| --- | --- | --- |
| Spatial associative learning | Water maze | Normal<sup>23</sup> but less precise/efficient searching<sup>21</sup> |
|  | Plus-maze | Normal<sup>22</sup> |
|  | Barnes maze | Normal, but more errors during training<sup>47</sup> |
|  | Place preference (IntelliCage) | Normal<sup>23</sup> |
| Spatial reversal learning | Water maze | Impaired (delayed acquisition/higher perseverance)<sup>21</sup> |
| Spatial reversal learning | Plus-maze | Normal<sup>22</sup> |
| Spatial rule-shift learning | Plus-maze (ego→allocentric) | Impaired (delayed acquisition, but not due to perseverative errors)<sup>22</sup> |
| Aversive associative learning | Cue FC | Normal<sup>23,24</sup>, or enhanced freezing in novel context<sup>24</sup>, or reduced freezing in novel context after repeated cue presentation<sup>18</sup> |
|  | Context FC | Normal<sup>23,24</sup> or reduced<sup>18</sup> freezing |
|  | Trace FC | Improved<sup>23</sup> |
| Operant learning | Nose-poking for reward | Normal<sup>23</sup> |
| Procedural learning | (various) | Normal<sup>23</sup> |
| Anhedonia | Sucrose preference | Normal, but transient anhedonia (delayed preference)<sup>24</sup> |
| Behavioural despair | Forced swim test, Tail suspension test | Mildly anti-depressive (reduced immobility)<sup>48</sup> |
| Nesting | Nest building | Reduced<sup>24</sup>, potentially due to avoidance of novel nesting material<sup>18</sup> |
| Sensitivity to psychostimulant | Amphetamine-induced locomotion | Enhanced<sup>18</sup> |
| Stereotypy | Stereotypic movements | Decreased<sup>24</sup> |
| Processing of novelty | Novelty-induced locomotion | Enhanced<sup>24</sup> or normal<sup>23</sup> |
|  | Novelty-induced exploration | Enhanced<sup>24</sup> |
|  | Exploration of novel material, incl. bedding | Avoidance of novel material<sup>18</sup> |
|  | Digging and marble burying | Reduced<sup>24</sup> |
| Unconditioned anxiety | Elevated Plus-maze | Normal<sup>23</sup> |
|  | Open field | Normal<sup>23</sup> |
| Olfaction | (Various) | Normal<sup>18</sup> or reduced<sup>23</sup> |
| Vision, tactile sense | (Various) | Normal<sup>18</sup> |
| Posture, gait, muscle tone | (Various) | Normal<sup>18</sup> |

Importantly, the same mouse line also shows a lack of neurogenesis in the adult dentate gyrus ^23^, which adds to its translational value given that indicators of reduced neurogenesis were also found in post-mortem brains from patients with schizophrenia ^25^. Furthermore, PV-cell hypofunction could be contributing to the neurogenesis deficit, since the innervation of new-born cells by PV-interneurons is critical for their survival and development ^26^. In reverse, new-born granule cells activate strong dentate gyrus-feedback- and CA3-feedforward inhibition ^27,28^, implying that reduced neurogenesis *alone* might lead to hippocampal hyperactivity. Importantly though, physiological and some behavioural aberrations in the CD2-KO mouse, could be rescued by implantation of GABAergic precursor cells specifically into the ventral hippocampus, pin-pointing this part of the hippocampus and its over-activity as the primary cause of those abnormalities ^18^. This suggests that the *Ccnd2^−/−^* mouse model might be well-suited to study the role of ventral hippocampal hyperactivity in the causation of schizophrenia-related deficits. To that end, we have subjected *Ccnd2^−/−^* mice to a large battery of behavioural tests in the positive, negative, and cognitive symptom domains.

## METHODS

### Animals

*Cyclin-D2* knockout *(Ccnd2^−/−^*, CD2-KO; 18 males, 12 females) mice ^17^ and wild-type littermate controls (19 males, 9 females) were generated by crossing heterozygous *Ccnd2^-/+^* mice. Animals were group-housed in Type II-Long individually ventilated cages (Greenline, Tecniplast, G), enriched with sawdust, Sizzle nest and cardboard houses (Datesand, UK), and subjected to a 13 h light / 11 h dark cycle. All experiments conformed to the German Animal Rights Law (Tierschutzgesetz) 2013 and were approved by the Federal Ethical Review Committee (Regierungsprädisium Tübingen) of Baden-Württemberg.

### Behavioural tests

Detailed protocols for all behavioural tests are given in Supplementary Methods and the majority of protocols were as previously described ^29^. Light levels were set to 100 lux for all tasks in open arenas. In brief, in the *novel-object recognition test*, mice were exposed to two identical objects, A1 and A2, in the sample phase (10 min) and – after a delay of 2 min – to an identical copy of the familiar object, A3, and a novel object, B1, in the test phase (5 min).

Direct physical interaction between the snout or whiskers with the object were scored manually by an observer blind to genotype, and the extent of novel-object recognition was determined according the ratio of interaction with B1 to interaction with B1 and A3 combined. *Spatial novelty preference testing* in the Y-maze was conducted analogously: animals explored the start arm and a pseudo-randomly selected goal arm in the sample phase (5 min), while they had access to all arms (including the previously unvisited novel arm) in the test phase (2 min) that followed after a 1 min-delay period during which the sawdust in the maze was mixed to remove olfactory landmarks. In the *T-maze test of working memory*, 10 trials were conducted per day in food-deprived (85–90 % of free-feeding weight) animals. Each trial consisted of a sample run with a forced choice into one of the goal arms, and a choice run in which the previously unvisited arm was baited and its visit was counted as a correct choice. Animals were trained up over 5 days in a blocked paradigm with a delay between sample and choice run of 5 s and an inter-trial interval (ITI) of 3–10 min. Subsequently, the protocol was altered every second day starting with a massed design with a delay of 1 s and an ITI of 25 s, followed by protocols in a blocked design with delays of 5, 20, or 60 s. The assessment of *spatial reference learning, rule-shift and rule-reversal* were conducted in a plus-maze with transparent walls where the start arm (North or South) was chosen pseudo-randomly and one of the goal arms (East or West) was baited according to the respective rule and its visit counted as a correct choice. 10 trials were given per day with a delay of 5–10 min between them. Animals were first trained up on an egocentric rule *(turn left* or *turn right*). After achieving the performance criterion of 85 % correct choices in 20 consecutive trials, animals were switched to an allo-centric rule (*go West* or *go East*) to assess spatial rule-shift learning. After reaching the same 85 % performance criterion the allo-centric rule was reversed (e.g. from *go West* to *go East*, or vice versa) to assess reversal learning.

*Unconditioned anxiety* was measured on the elevated plus-maze, according to the preference with which animals explored the two open vs the two closed arms. *Novelty-induced locomotion* was assessed by letting animals explore a new Typ-III-cage (Tecniplast, G) with clean sawdust for 2 h. Movement was video-tracked using ANY-maze (San Diego Instruments, CA, USA) in the Y-maze, the elevated plus-maze and during novelty-induced locomotion. *Reciprocal social interaction* was assessed in similar cages but with darker walls (Tecniplast), into which a younger adult stimulus mouse of the same sex and strain had been introduced. Social interactions (sniffing, taxing with snout or whiskers, but not aggression or sexual behaviour) were scored offline manually and blind to genotype over a period of 16 min in 2 min intervals.

*Sucrose preference* was assessed overnight in animals that were single-housed on the first day of the testing sequence, on which they were presented with two bottles of drinking water. Over the 4 subsequent nights one of the two bottles contained either 1 % w/v (2 nights) or 10 % w/v (2 nights) sucrose in drinking water, while only the water bottle remained in the cages during the day and for the sixth night. During this last night, animals received a Nestlet™ (Datesand) of 2.3–2.6 g, and nests were scored according to an established *nest building* rating system ^30^, next morning.

For the *5-choice-serial reaction time task* (5-CSRTT) mice were trained up in operant chambers (Med Associates Inc., VT, USA) as previously described ^29^ over 4 stages with increasing attentional difficulty until mice reached a consistent performance (≥80 % accuracy, ≤50 % omissions, ≥30 trials) on the stage 4 baseline protocol where the stimulus duration was 4 s and the inter-trial interval (waiting) time was 5 s. Subsequently, various protocols to challenge either *sustained attention* (by reduction of the stimulus duration to 1 s or distraction, or both) or *impulse control* (by extension of the waiting time to 9 s) were applied over 2 consecutive days, followed by 1–3 d of baseline training before the next challenge. Afterwards, mice were trained on a simplified version of this task where only hole 2 or 4 was illuminated in each trial until mice showed an accuracy of ≥80 % in two consecutive sessions. Subsequently, the rule was shifted from a cued to a spatial version, whereby only hole 2 (or, in other mice, hole 4) was rewarded irrespective if hole 2 or 4 was illuminated in a given trial. To assess *rule-shift learning* in this operant paradigm, mice were trained until they had reached a criterion of 80 % accuracy on the trials in which the correct hole was unlit on 3 consecutive days. Condensed milk diluted with drinking water was used as reward on rewarded maze tasks, strawberry milk was used in operant box paradigms.

### Behavioural pharmacology

To assess the effects of the D1-antagonist SCH23390, the D2-antagonist raclopride and the mGlu2/3R-agonist LY379268 on *novelty-induced locomotion* in CD2-KO mice, each drug was injected i.p. at an injection volume of 10 ml/kg prior to an 1 h exposure in a novel open-field, identical to those used for the prior assessment of novelty-induced locomotion (see above). S(-)Raclopride-(+)tartrate salt (Sigma, G) was diluted in 0.3 % TWEEN80/saline and injected 60 min before the start of the run. SCH23390 hydrochloride and LY379268 disodium salt (Tocris, G) were diluted in saline and injected 30 min or 45 min, respectively, before the run. For *dose-response assessments* of these drugs, separate adult male C57BL/6 cohorts were used in a between-subject design (4–8 mice per sub-group); either raclopride, SCH23390 or LY379268 were injected 30 min, 30 min, or 45 min, respectively, before a 30 min exposure to the novel open-field, followed by injection of 0.2 mg/kg MK-801 maleate (in saline) and another 60 min of locomotor monitoring. For the *T-maze test*, LY379268 was injected 50–60 min before testing commenced. Additionally, amphetamine-induced locomotion was tested by injection of D-amphetamine hemisulfate (Sigma, diluted in saline), 30 min after an acclimation period in the novel open-field, immediately followed by another hour of post-injection exposure. Latin-square designs with a fixed decreasing order of doses, counterbalanced for starting dose and genotype, were used for all within-subject studies.

### Analysis

Within-subject data was analysed by repeated-measures ANOVA, and between-subject data with univariate ANOVA (subsequently termed “ANOVA” only), one-way ANOVA, *t*-Tests, or - in cases of non-parametric pairwise comparisons - the Mann-Withney-U-Test (termed MWU), as appropriate. All graphs display mean ± S.E.M.

## RESULTS

### Sustained novelty-induced hyperlocomotion in CD2-KO mice

Novelty-induced hyperlocomotion was assessed over a 2 h exploration time. A repeated-measures ANOVA over this period split into 24 5-minute intervals revealed a significant effect of genotype (*P* < 0.0005) and a trend for gender (*P* = 0.051), but no interaction between the two factors (*P* = 0.791), indicating that KO mice are hyperlocomotive (Figure 1a, b; see Supplementary Table S1 for statistical details on this and all subsequent results including n-, F- and t-values). Additionally, the number of rotations, an indicator of stereotypies, was significantly elevated in knockouts of both genders (*P*_genotype_ = 0.001; *P*_gender_ = 0.237, *P*_genotype*gender_ = 0.932; ANOVA, Figure 1c).

**Figure 1.**
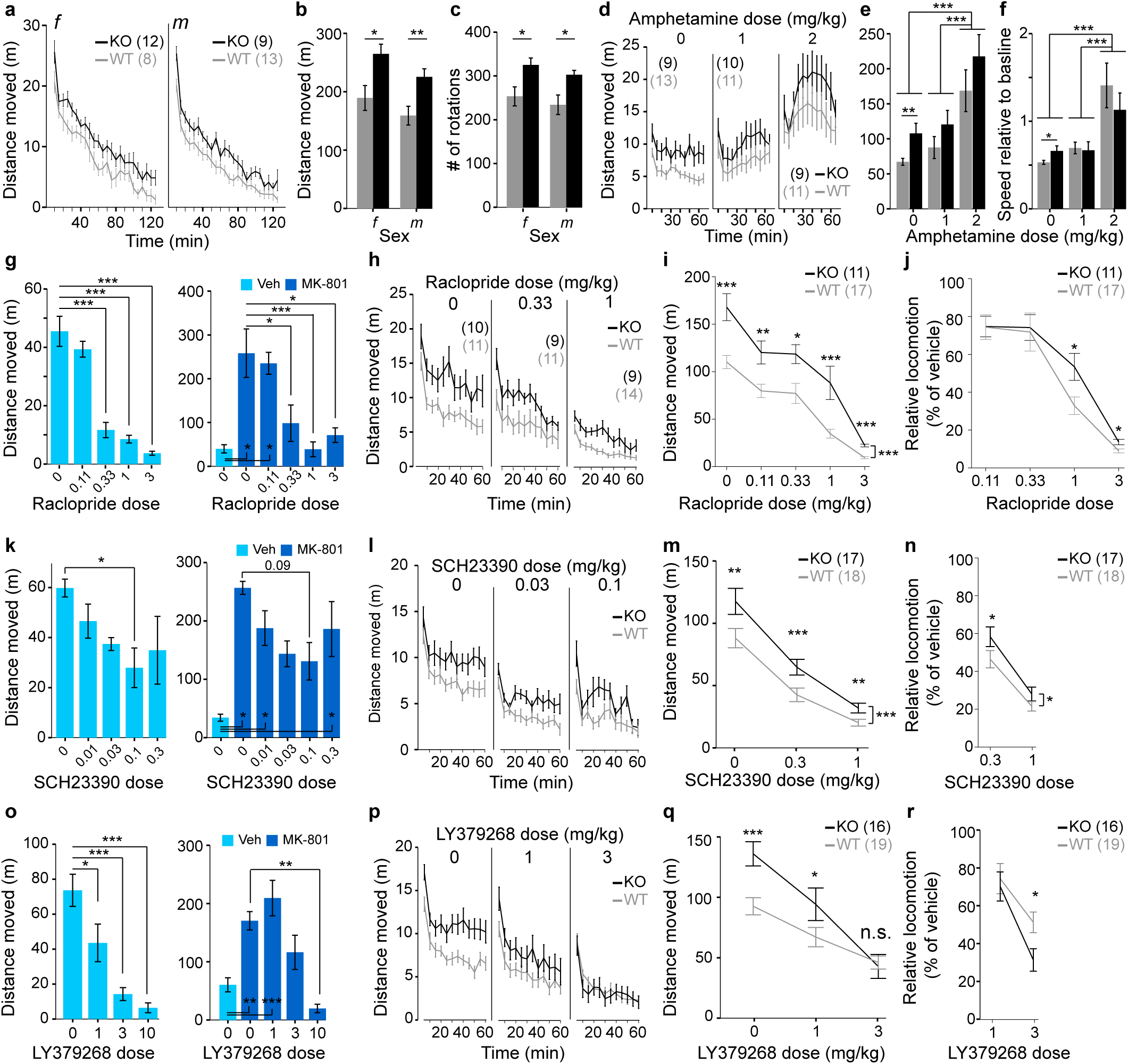
Pharmacology of novelty-induced hyperlocomotion in CD2-KO mice. **(a, b)** Locomotion in a novel environment in 5 min intervals (a) and total over 2 h (**b**). (**c**) Number of rotations in the same experiment. **(d, e)** Locomotion after injection of vehicle (0), 1 or 2 mg/kg amphetamine in 5 min intervals **(d)** and in total over 2 h **(e)**. **(f)** Same data as in **(e)** but expressed as speed (m/min) relative to the speed in the 30 min preceding the amphetamine injection. (**g, k, o**) Dose-response measurements on the efficacy of raclopride **(g)**, SCH23390 **(k)** and LY379268 **(o)** in reducing novelty-induced (left) and MK-801-induced (right, dark-blue indicates injections of 0.2 mg/kg MK-801, light-blue is vehicle) hyperlocomotion using the stated doses (x-axes). **(h)** Between-subject assessment of the efficacy of raclopride at the stated doses on novelty-induced locomotor activity in CD2-mice in 5 min intervals (total distance moved and statistics not shown). (i, m, q) Within-subject assessment of raclopride **(i)**, SCH23390 **(m)** and LY379268 **(q)** at the stated doses on novelty-induced locomotion, plotting total distance moved in 1 h. (**j, n, r**) Within-subject assessment of raclopride **(j)**, SCH23390 **(n)** and LY379268 **(r)** at the stated doses on novelty-induced locomotion, plotting distance moved in 1 h (as %) relative to the vehicle (0) value in each animal. **(l, p)** Same data as in **(m, q)**, within-subject assessment of SCH23390 **(l)** and LY379268 **(p)** at the stated doses on novelty-induced locomotion, plotting distance moved in 5 min intervals. Wildtype controls, grey; CD2-KO, black; *f*, females, m, males; numbers in brackets denote *n* for each group. All doses stated in mg/kg. * *P* < 0.05, ** *P* < 0.01, *** *P* ≤ 0.001. For trends, *P* values are stated as numbers. Error bars, S.E.M..

### Normal amphetamine-induced hyperlocomotion in CD2-KO mice

In a similar locomotor activity paradigm, injection of 2 mg/kg amphetamine after an acclimation period of 30 min resulted in a significant subsequent increase of locomotion relative to vehicle and 1 mg/kg amphetamine conditions in both genotypes *(P*_dose_ < 0.0005, *P*_genotype_ = 0.015, *P*_genotype*dose_ = 0.957, ANOVA; between-subject design; Figure 1d-e). Univariate ANOVA within each dose revealed that the higher locomotion of CD2-knockouts relative to controls seen qualitatively at each dose, only reached significance after vehicle injection (*P* = 0.007) but not after amphetamine injections (*P* > 0.1). Likewise, when expressing locomotor activity post-amphetamine relative to locomotor activity before injection there was no difference between genotypes (*P* = 0.759), while the effect of dose remained (*P* < 0.0005, ANOVA). This suggests, that CD2-KO mice are not more sensitive to amphetamine than controls; if anything, amphetamine aligns wildtype with knockout levels of locomotor activity (Figure 1f).

### Incomplete reduction of novelty-induced locomotion by the D2/3 antagonist raclopride

Signalling through D2-type receptors in the nucleus accumbens mediates spontaneous locomotor activity in rodents ^31^, and antagonism of these receptors is also the suggested mechanism of the antipsychotic action of neuroleptics. We therefore tested, if the novelty-induced hyperlocomotion in CD2-KO mice is indeed responsive to the D2-receptor antagonist raclopride. We first confirmed in a dose-response study in a separate group of male C57BL/6J mice, that the doses 0.33, 1 and 3 mg/kg raclopride were effective in reducing novelty and MK-801-induced hyperlocomotion, while 0.11 mg/kg was statistically ineffective (Figure 1g). Using the two lowest effective doses 0.33 and 1 mg/kg, we firstly performed a between-subject study in CD2-KO mice and controls. While raclopride at both doses significantly reduced locomotion (distance moved) in both genotypes, surprisingly, knockout mice remained significantly hyperlocomotive at both doses compared to wildtype controls (*P* < 0.0005, ANOVA; *P* < 0.05 for ANOVAs at each dose, Figure 1h). That means, although 0.33 and 1 mg/kg could reduce locomotion in knockouts to the level of vehicle-injected controls, these doses could not reduce locomotion to control levels *within* each dose. Also, a dose of 0.33 mg/kg could significantly reduce the number of rotations in wildtype mice compared to vehicle (*P* = 0.004, Tukey post-hoc test), but not in knockout mice (*P* = 0.646), suggesting that the drug is even less effective in the latter (Supplementary Table S1).

To investigate this further, a subset of mice were continued in a within-subject regime with injections of vehicle and the doses 0.11, 0.33, 1 and 3 mg/kg. Again, at all doses, knockout mice remained hyperlocomotive compared to wildtype controls (*P* < 0.0005, repeated-measures ANOVA; *P* < 0.05 for univariate ANOVAs). Across all 5 doses a significant dose*genotype interaction was apparent (*P* = 0.033, Figure 1i), indicating a differential efficacy of raclopride. Expressing the amount of locomotion as percentage of the locomotion each mouse displayed in its vehicle-trial revealed that the higher doses, 1 and 3 mg/kg, reduced novelty-induced locomotion significantly *less* in knockout mice compared to wildtype controls (*P* < 0.05, ANOVA, Figure 1j), supporting the view that the drug is less effective in CD2 knockouts. This suggests, that while D2-receptors may mediate baseline locomotion which is reduced in both genotypes, there is an additional, D2-independent mechanism of increasing locomotor drive in CD2 knockouts. This is reminiscent of the hyperlocomotion produced by electrical (20 Hz) stimulation of the ventral hippocampus in rats, which is mediated by D1- rather than D2-receptors ^5^.

### Incomplete reduction of novelty-induced locomotion by the D1 antagonist SCH23390

Therefore, we repeated this experiment with the D1-antagonist SCH23390. Assessing the efficacy of 0.011, 0.033, 0.1 and 0.3 mg/kg against novelty- and MK-801-induced locomotion in a separate cohort of C57BL6/J males revealed that the two intermediate doses 0.033 and 0.1 mg/kg had the strongest, yet statistically moderate, effects (between-subject design; Figure 1k). Those two doses were tested against novelty-induced locomotion in CD2-KO mice and controls (within-subject design). While the drug significantly reduced locomotion dose-dependently in both genotypes, KO mice again remained hyperlocomotive relative to wildtype controls receiving the same dose (*P*_genotype_ < 0.0005, *P*_dose_ < 0.0005, *P*_dose*genotype_ = 0.016, repeated-measures ANOVA; Figure 1l-m). Expressing locomotor activity under each dose relative to distance moved after injection of vehicle revealed that the drug was less effective in reducing hyperlocomotion in CD2-KO mice compared to wildtypes *(P*_genotype_ = 0.02, repeated-measures ANOVA; Figure 1n).

### Normalization of novelty-induced locomotion by the mGlu2/3 agonist LY379268

mGlu2/3 agonists have been proposed to be able to scale down glutamatergic transmission and one compound from this family, LY379268, has been shown to normalize ventral hippocampal hyperactivity evoked by chronic ketamine application in mice ^2^. In a separate cohort of male C57BL6/J mice, we found that a dose of 1 mg/kg LY379268 could reduce novelty-induced but not MK-801-induced hyperlocomotion, while 3 mg/kg strongly reduced novelty-induced locomotion and moderately normalized MK-801-induced locomotion (Figure 1o). Testing both doses in CD2-KO mice (within-subject design), we found that 3 mg/kg can completely normalize novelty-induced locomotion in CD2-KO mice relative to wildtype controls receiving the same dose (*P*_genotype_ = 0.941), while 1 mg/kg achieved a partial reduction (*P*_genotype_ = 0.029 at 1 mg/kg and *P*_genotype_ < 0.0005 at vehicle, ANOVA, Figure 1p-q). Indeed, LY379268 appeared to reduce locomotion even stronger in knockouts than in wildtypes (*P*_genotype*dose_ = 0.01) leading to significantly less locomotion relative to the vehicle trial at 3 mg/kg compared to wildtype controls (*P*_genotype_ = 0.043, ANOVA; Figure 1r). We confirmed the finding of a significant interaction between dose (vehicle vs 3 mg/kg) and genotype in a between-subject study in an extended cohort (*P*_genotype*dose_ = 0.015, n: 15 KO, 20 WT; not shown).

### CD2-KO mice are impaired in various forms of cognitive flexibility

Impairments of executive function are a hallmark of schizophrenia and reduced capacity to learn a rule-shift in a spatial version of the attentional set-shifting task conducted in the plus-maze has been demonstrated in *Ccnd2^−/−^* mice previously ^22^. Firstly, we sought to confirm this finding. KO mice acquired a spatial long-term memory according to an egocentric rule similarly well as wildtypes. The number of trials needed to reach the criterion of 85 % correct of the last consecutive 20 trials did not differ significantly between genotypes (*P* = 0.232, *t*-test), and the learning curves were similar (*P*_genotype_ = 0.367, repeated-measures ANOVA over the second half of the 12 training blocks conducted by all animals; Figure 2a-c). This indicates, that CD2-KO mice can process spatial information and acquire a spatial memory largely normally. Once animals had reached criterion, the associative rule was shifted to an allo-centric one ^22^. In this task, KO mice needed significantly more trials to reach criterion again (*P* = 0.04, *t*-test) and average performance was lower compared to wildtypes (*P* = 0.048, repeated-measures ANOVA over the second half of the 14 training blocks conducted by all animals; Figure 2d-f).

**Figure 2.**
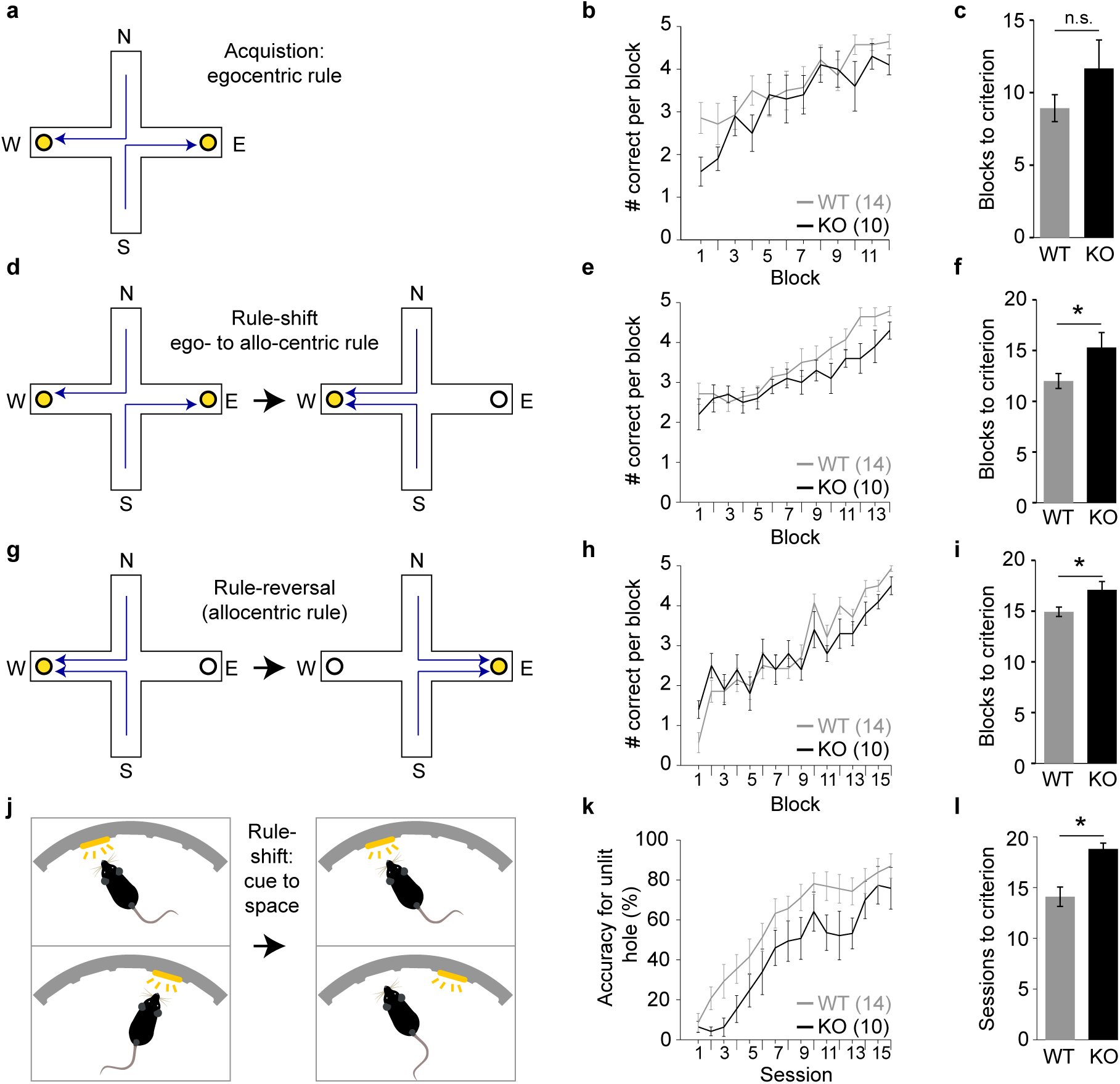
Impairments of cognitive flexibility in CD2-KO mice. (**a, d, g**) Sequential learning paradigms in the plus-maze starting with the acquisition of an egocentric appetitive spatial reference memory **(a)**, followed by a rule-shift towards an allocentric associative memory **(d)**, and a rule-reversal within the allocentric rule **(g)**; specific rules (arrows) are shown as examples only. Goal arms are denoted West **(W)** and East **(E)**, start arms North **(N)** and South **(N)**. (**b, e, h**) Number of correct choices in blocks of 5 trials for the number of blocks that were done by all mice in the cohort in the reference learning **(b)**, the rule-shift learning **(e)** and the rule-reversal learning **(h)**. (**c, f, i**) Blocks of 5 trials needed to reach the criterion of 85 % in the last 4 blocks (20 trials) correct choices in the reference learning **(c)**, the rule-shift learning **(f)** and the rule-reversal learning **(i)**. **(j)** Operant paradigm of rule-shift learning from a hole indicated by the cue-light to a hole defined by its spatial position (while cue lights are still presented but bare no prediction). **(k)** Accuracy for the correct choice in trials where the incorrect hole is illuminated in each session. **(l)** Number of sessions needed to reach the criterion of ≥80 % accuracy in three consecutive sessions. Wildtype controls, grey; CD2-KO, black. * *P* < 0.05. Error bars, S.E.M..

Once mice had reached criterion again, rule-reversal learning was assessed: the allo-centric spatial location that was associated with a reward was switched. Similarly to the rule-shift, KO mice required more trials than controls to reach criterion again (*P* = 0.021, *t*-test) and their performance was inferior (*P* = 0.01, repeated-measures ANOVA over the second half of the 16 training blocks conducted by all mice).

Given that rule-shift learning is usually attributed to prefrontal rather than hippocampal dysfunction, we sought confirmation of our surprising finding by conducting a newly developed operant-box paradigm that resembles the Wisconsin card-sorting test (which is indicative of prefrontal dysfunction in humans) by implementing a rule-shift within the visual domain (Figure 2j, also see Methods): mice that had been trained up in the 5-choice-serial-reaction-time task (see below) in which a rewarded nose-poke is indicated by a cue light, were subjected to a rule-shift whereby cue-lights were rendered meaningless and the spatial location now indicated the correct choice. KO mice required significantly more trials to reach the criterion of 80 % accuracy in the choice of the unlit rewarded hole on three consecutive days (*P* = 0.006, ANOVA) and showed lower accuracy during training (*P* = 0.006, repeated measures ANOVA over the second half of the 16 training blocks conducted by all mice, Figure 2k-l; 1 wildtype and 2 CD2-KO mice were excluded as they did not reach criterion within 23 sessions).

### CD2-KO mice are unimpaired in object-related and spatial short-term habituation

Dysfunction of sustained and stimulus-specific attention (attentional control) have been related to schizophrenia ^32,33^, and could explain deficits in other cognitive tests conducted in this study. We first assessed a form of spatial short-term memory that is largely mediated by short-term habituation of attention using the spatial novelty-preference test in the Y-maze ^34^. In this test, habituation is measured as a preference for a novel as opposed to a familiar choice arm (or ‘space’). In the sample phase, KO mice showed significant hyperlocomotion (*P* = 0.018, univariate ANOVA) but spent a similar amount of time in the to-be-familiar arm (*P* = 0.425) as wildtype mice suggesting equal levels of exploration (Figure 3a-c). In the test phase, mice of both genotypes explored the novel arm significantly more than the familiar arm, in terms of both entries and time (*P* < 0.0005, repeated-measures ANOVA, Figure 3d-e), and there were no genotype-related differences in the preference ratios for the novel arm (*P* > 0.2, ANOVA; Figure 3f). Additionally, preference ratios were significantly above chance level in both genotypes (*P* < 0.005, one-sample *t*-test).

**Figure 3.**
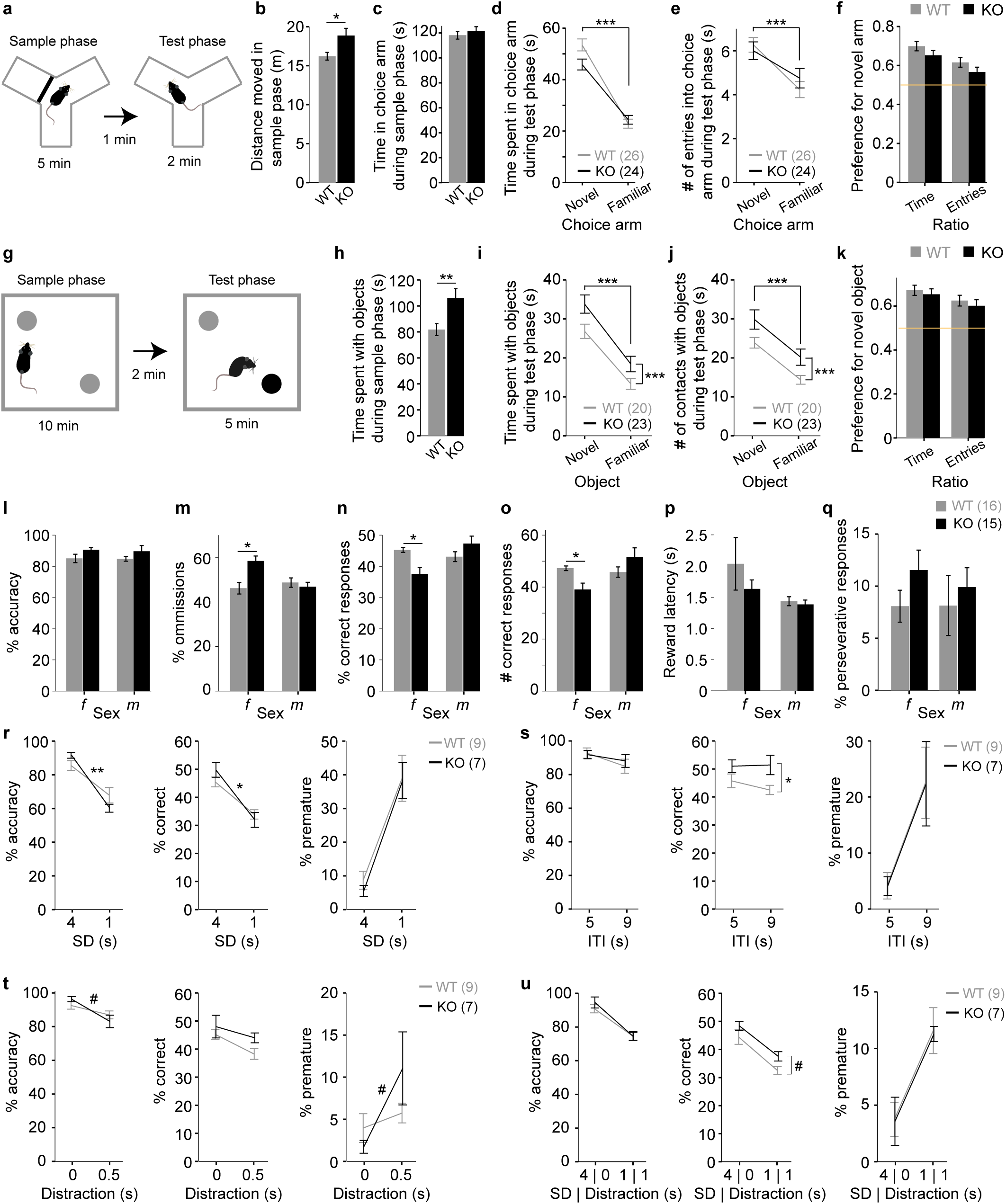
Intact sustained attention and short-term habituation in CD2-KO mice. **(a)** Paradigm of the spatial-novelty preference test. **(b)** The total distance moved and **(c)** time spent in the to-be-familiar choice arm in the sample phase. **(d)** Time spent in the indicated choice arms and **(e)** number of entries into them during the test phase. **(f)** Preference for the novel arm according to either time or entry measures. **(g)** Paradigm of the novel-object recognition task, different colours indicate different types of objects. **(h)** Amount of time the animals investigated both objects in the sample phase. **(i, j)** Interaction with the novel and the familiar object in the test phase in terms of time **(i)** and number of contacts **(j)**. **(k)** Preference for the novel object according to either time or entry measures. (**l-q**) Performance in measures of attention (accuracy **(l)**, percent omissions **(m)**, percent **(n)** and number of correct responses **(o)**), incentive motivation (latency to reward receptacle **(p)** and perseveration (percent perseverative responses into the correct hole, **(q)**) in the baseline protocol (4 s stimulus duration, SD; 5 s inter-trial-interval, ITI; average of the last 4 sessions before the challenges started). (**r-u**) Measures of attention (accuracy, left; %correct, middle) and impulsivity (percent premature responses relative to all initiated trials, right) various challenge protocols (right value in all subpanels, average of 2 consecutive sessions) relative to the baseline protocol immediately before the challenge (left value in all subpanels). Attention was challenged by a reduction of the SD to 1 s **(r)**, a random distraction by switching of the house light for 0.5 s during the ITI **(t)** and a combination of SD-reduction to 1 s and random switching of the house light for 1 s **(u)**. Impulsivity was challenged by increasing the waiting time (ITI) until the holes were illuminated to 9 s **(s)**. Wildtype controls, grey; CD2-KO, black; *f*, females, m, males; numbers in brackets denote *n* for each group. ^#^ *P* < 0.1, * *P* < 0.05, ** *P* < 0.01, *** *P* ≤ 0.001. In within-subject data (line graphs), symbols on horizontal lines indicate an effect of the within-subject factor (not drawn in (r-u) for clarity), symbols on vertical lines indicate effects of genotype, and symbols on the connecting lines indicate an interaction between both factors. The yellow lines indicate chance-level performance.Error bars, S.E.M..

Novel-object recognition represents an object-related version of the novelty-preference test of habituation-driven short-term memory (Figure 3g) ^33,35^. CD2-KO mice, compared to controls, showed strongly increased exploration of the objects in the sample phase and the test phase (*P* < 0.01; ANOVA; Figure h-j). However, mice of both genotypes showed significantly higher exploration of the novel compared to the familiar object in the test phase (*P* < 0.0005, repeated-measures ANOVA), and did not differ in their preference for the novel object (*P* > 0.4, ANOVA; Figure 3k). In summary, this indicates, that CD2-KO mice are not impaired in short-term habituation of attention.

### CD2-KO mice are not impaired in sustained attention, motivation, perseveration or impulse control

To assess sustained attention, CD2-knockouts and wildtype controls were trained on the 5- choice serial reaction time task (5-CSRTT) through 4 stages of increasing difficulty, before being challenged with different protocols (see Supplementary Methods). In the baseline protocol, which was conducted with a stimulus duration (SD) of 4 s and an inter-trial-interval (ITI) of 5 s, knockout mice appeared significantly better in the primary measure of attention, % accuracy (*P* = 0.038; ANOVA), although in each of the gender groups alone there was no significant effect of genotype (*P* > 0.05, t-test; Figure 3l). In the secondary measures of attention, % omissions, % correct and number correct responses, female knockouts performed worse than female wildtype mice (*P* < 0.05, t-test), although, qualitatively, the opposite was observable in the male cohort, yielding significant genotype * gender interactions in all three measures (*P* < 0.02, ANOVA; Figure 3m-o). The basic measure of motivation, reward latency, did not differ between genotypes (*P* > 0.3, ANOVA), but females, irrespective of genotype, had a longer reward latency than males *(P_gender_* < 0.05, *P_gender*genotype_* = 0.307, ANOVA; Figure 3p). Additionally, perseveration, indicated by unrewarded repetition of nose-pokes into previously rewarded holes, was not significantly higher in CD2-KO mice compared to controls (*P* = 0.303, ANOVA; Figure 3q).

After baseline training, four challenge protocols were conducted, out of which three were targeted at sustained attention (shortening of the stimulus duration, distraction during the ITI, or both), and one at impulse control (increasing the ITI to 9 s). Within 60 sessions, 2 wildtype males and 7 of the 13 involved females (KO and WT) did not reach the criterion in the baseline stage to proceed to challenge protocols, which were therefore only conducted in males. Each challenge protocol was conducted over two consecutive days and was preceded and succeeded by sessions with the baseline protocol. For analysis of each parameter, the values from the two challenge days were averaged and a repeated-measures ANOVA was calculated by including the performance of the preceding baseline session.

Regarding impulse control, all challenges increased the primary measure of motor impulsivity, % premature responses (*P* < 0.0005 for all challenges except the exclusive distraction (*P* = 0.015), repeated-measures ANOVA). Similarly, all challenges decreased attentional accuracy (*P* < 0.0005 for all attention challenges, *P* = 0.001 for the impulsivity challenge). A trend of slightly better attention in CD2-KO mice at baseline (see above) was seen in the baseline sessions of most protocols (Figure 3r-u). However, in knockout mice attentional accuracy was affected significantly stronger by the first (and therefore unexpected) attentional challenge (*P*_challenge*genotype_ < 0.05 for %accuracy and %correct), and a similar trend (*P*_challenge*genotype_ = 0.066 for %accuracy) was seen with the first distraction challenge (Figure 3r, t). However, that effect was absent at the later combined SD-reduction/distraction challenge (Figure 3u), suggesting it does not reflect a genuine attentional deficit. With respect to %premature, there was no significant effect of genotype nor a genotype*challenge interaction (*P* > 0.05) detectable on any of the protocols (Figure 3r-u).

In summary, CD2-knockout mice appeared largely normal on all psychiatrically relevant parameters measured by the 5-CSRTT and were unimpaired in acquiring an associative memory during operant learning. While knockouts were unaltered in measures of motivation, perseverance and impulsivity, they showed slightly improved sustained attention at baseline, which was then, however, somewhat stronger disrupted by unexpected challenges. This is in line with the notion that schizophrenia might be associated with high but inflexible (“sticky”) attention, rather than blunt inattentiveness.

### CD2-KO mice show a working memory impairment that is resistant against LY379268

To assess spatial working memory, mice were trained on a rewarded alternation (delayed-non-match-to-place) task on the T-maze over 5 days in a blocked paradigm with a delay (intra-trial interval) of 5 s and an inter-trial interval of >3 min. While female knockouts did not differ from controls in this initial training paradigm, male knockouts performed worse across those training sessions (females: *P_genotype_* = 0.497; males: *P_genotype_* = 0.005; both genders: *P_genotype_* = 0.013, *P_gender_* = 0.002, *P*_genotype*gender_ = 0.123, *P*_day_ = 0.001, repeated-measures ANOVAs; Figure 4a-c). Subsequently, mice were challenged by different alterations of the paradigm each performed in two consecutive sessions: firstly, the delay was maximally reduced (“1 s”) and the paradigm changed to massed trials with an inter-trial interval of 25 s, which had revealed subtle deficits in mice with PV-interneuron-specific NMDA receptor hypofunction previously ^29,36^. Then, blocked paradigms with increasing delay (5 s, 20 s, 60s) were conducted. Across paradigms and genders, knockout mice were impaired (*P* < 0.0005), and females preformed significantly better (*P* < 0.0005; repeated-measures ANOVA). When assessing individual challenges by univariate ANOVA across genders, all protocols except the 60s-delay revealed significantly reduced performance in knockouts (*P* < 0.05; Figure 4d). (The 60 s-protocol strongly reduced performance in wildtype mice, especially in males, preventing the detectability of an effect of genotype.) When analysing the genders individually, effects of genotype dropped below significance level (*P* > 0.05, *t*-test) in the 20 s protocol in females, as well as in the 5 s (*P* = 0.059) and the 60 s-protocol in males (Figure 4e).

**Figure 4.**
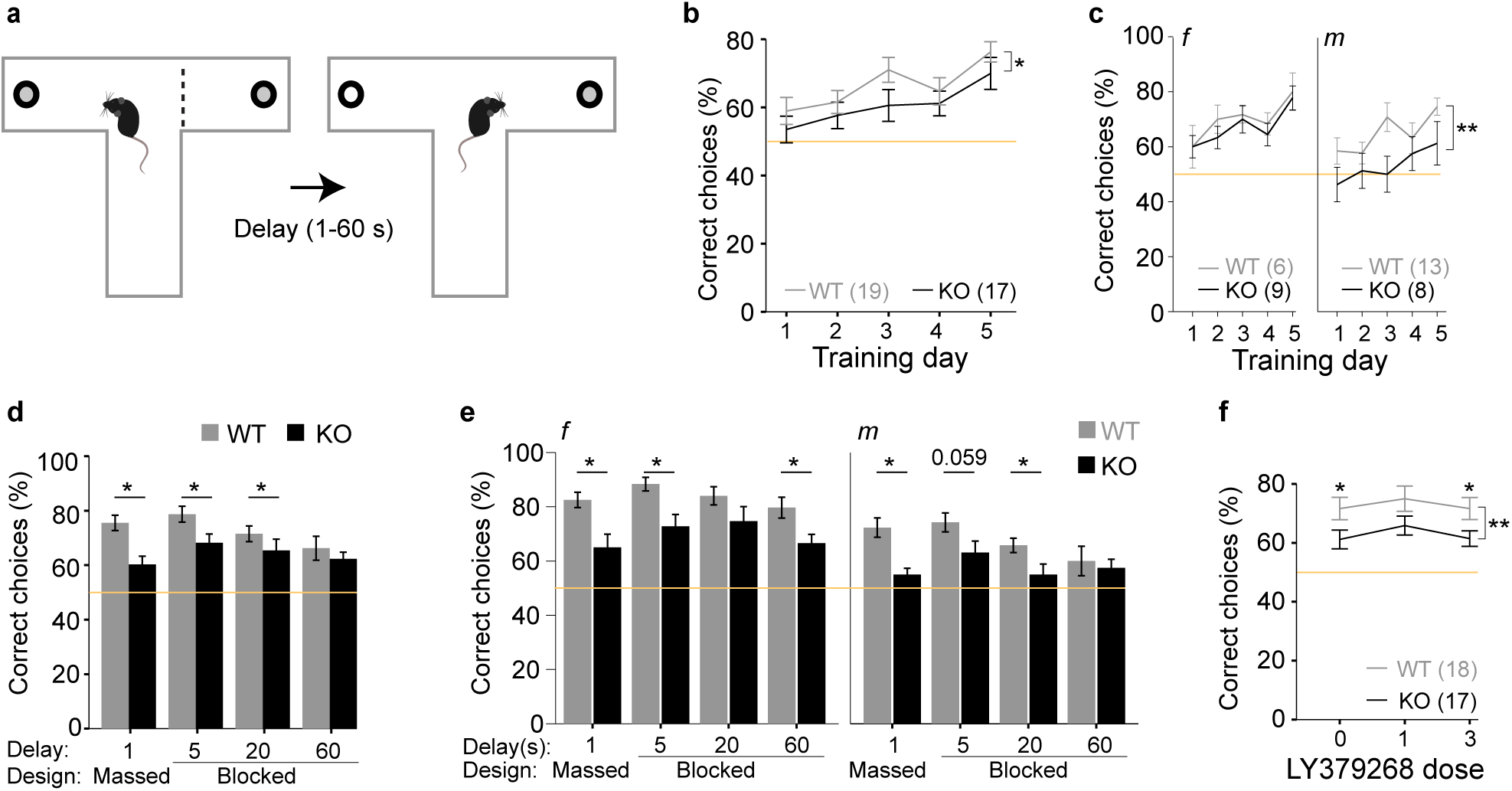
Impaired spatial working memory in CD2-KO mice. **(a)** Paradigm of the spatial working memory test. **(b, c)** Performance (correct choices out of 10 trials, in %) over the 5 initial training days shows a significant effect of genotype across the total cohort **(b)**, which is driven by the male subgroup (**c**, right), rather than the females (**c**, left). **(d, e)** Performance under 4 different challenge protocols conducted in the same order as plotted with 2 consecutive days per protocol (averaged for the display) shown for the whole cohort **(d)** and the genders separately **(e)**; protocols differed by their design (massed vs blocked) and the delay between sample and choice trial as indicated on the x-axes. **(f)** After a re-training (identical to the previous one **(b)**, data not shown) either vehicle (0), 1 or 3 mg/kg LY379268 was injected prior to testing sessions in the massed design (1 s delay), with interleaved re-training sessions (blocked design, 5 s delay, not shown). Wildtype controls, grey; CD2-KO, black; *f*, females, m, males; numbers in brackets denote *n* for each group. * *P* < 0.05, ** *P* < 0.01. In within-subject data (line graphs), symbols above data points indicate an effect of group in the indicated condition, symbols on vertical lines indicate effects of genotype across conditions. The yellow lines indicate chance-level performance. Error bars, S.E.M..

We next examined, if LY379268 was equally effective against the working memory deficit in CD2-KO mice as it was against their hyperlocomotion. However, across doses the lower performance of knockouts remained apparent (*P*_genotype_ = 0.007, *P* > 0.05 for effects of dose, gender and all interactions, repeated-measures ANOVA, within-subject design, Figure 4f).

### CD2-KO mice show normal unconditioned anxiety

During the testing of novelty-induced locomotion in an open field (see above), we found no effect of genotype on the average distance to the cage walls (*P* > 0.1 for all factors, ANOVA, Figure 5a), suggesting similar levels of unconditioned anxiety. Likewise, on the elevated plus-maze, animals of both genotypes spent significantly more time in the closed compared to the open arms (*P*_arm_ < 0.0005, repeated-measures ANOVA, Figure 5b). Accordingly, the preference ratio for the open arm (time spent in open arms divided by time spent in all arms) was not significantly different between genotypes (*P* = 0.488), but was significantly lower in females compared to males (*P* = 0.003, univariate ANOVA, Figure 5c). Preference ratios calculated from the *number* of entries did not differ between any groups (*P* > 0.1, ANOVA, Figure 5d). This confirms that anxiety is unaltered in CD2-knockouts.

**Figure 5.**
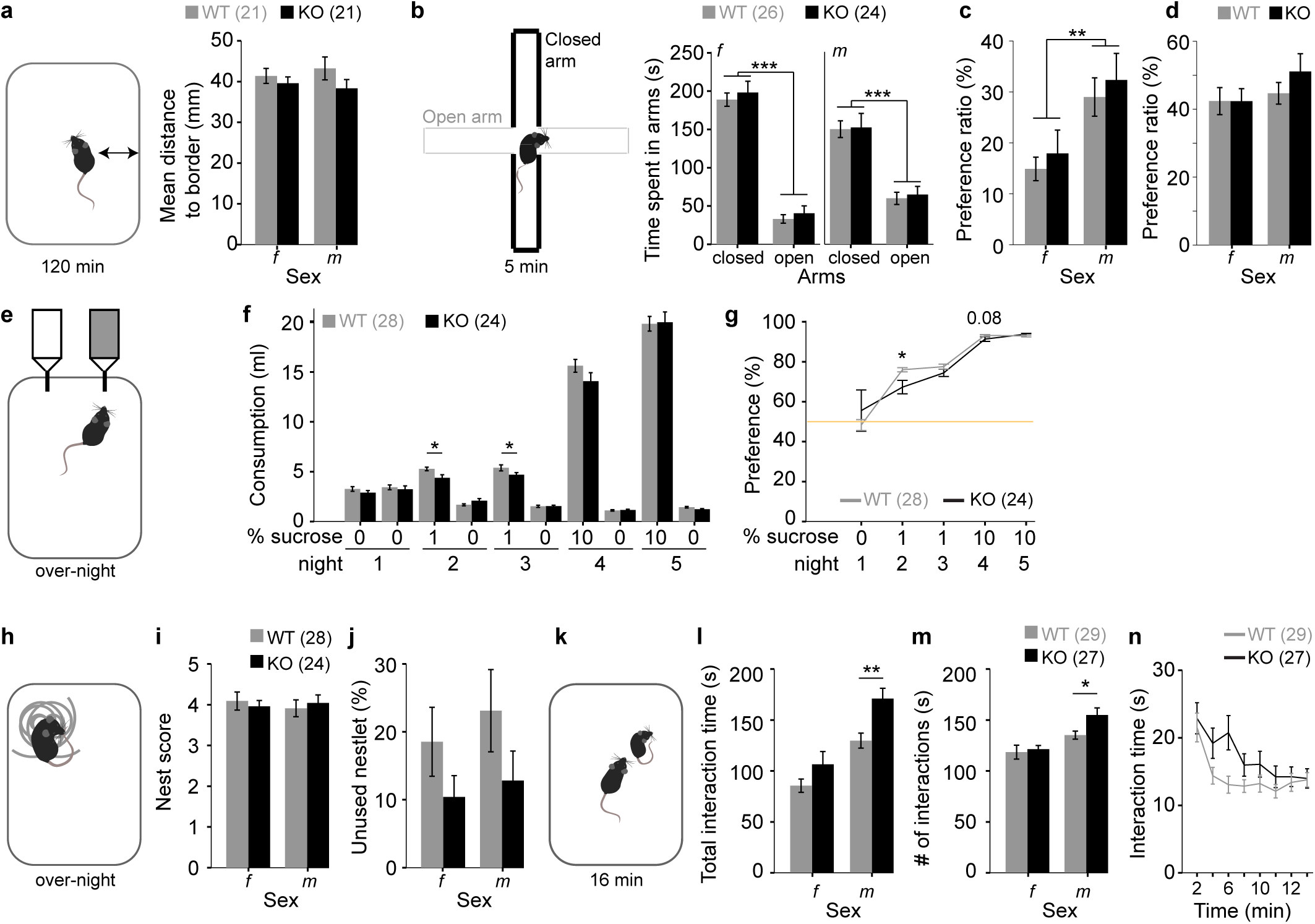
Intact affective functioning of CD2-KO mice. **(a)** Paradigm and data of measuring unconditioned anxiety according to the average distance from the wall of an open field during 2 h of exploration. (**b-d**) Paradigm and data of measuring unconditioned anxiety in the elevated plus-maze. **(b)** Time spent in the open and closed arms as indicated, and **(c)** the preference ratio calculated from those times (time spent in open arms/time spent in all arms). **(d)** Preference ratio calculated as in **(c)** but from the number of entries into open and closed arms. (**e-g**) Paradigm and data from the sucrose preference test of anhedonia. **(f)** Absolute consumption volumes from the bottles with indicated sucrose concentration in drinking water over 5 nights, and **(g)** preference ratios (consumed volume of sucrose-water/total consumed liquid) over the 5 nights, with the yellow line indicating lack of preference. Note that both bottles contained water in the first night. (**h-j**) Paradigm and data for nest-building assessment according to **(i)** the qualitative nest score (1–5 rating scale with 5 indicating the best nests ^30^), and **(j)** left-over untorn material relative to weight of the original nestlet. (**k-n**) Paradigm and data for assessment of reciprocal social interaction with an unknown younger adult mouse of the same sex and strain. **(l)** Total time of social interactions and **(m)** total number of interaction over 16 min. **(n)** Interaction time in 2 min bins. Wildtype controls, grey; CD2-KO, black; *f* females, m, males; numbers in brackets denote *n* for each group. * *P* < 0.05, ** *P* < 0.01, *** *P* ≤ 0.001. For trends, *P* values are stated as numbers. Error bars, S.E.M..

### CD2-KO mice show transient anhedonia in the sucrose preference test

Analysing the preference ratio for sucrose (consumed volume of liquid from sucrose bottle relative to total consumed liquid) over five nights, we found a strong effect of night (indicating sucrose concentration; *P* < 0.0005) but also a night*genotype interaction (*P* = 0.007) without an effect of genotype (*P* = 0.647, repeated-measures ANOVA), suggesting that both genotypes develop a strong preference for sucrose but with a different trajectory (Figure 5e-g). This divergence, resulted largely from the first encounter with sucrose (night 2), where an interaction between genotype and bottle-content (water vs sucrose, *P* = 0.001) on the consumed liquid volume, as well as a significantly reduced consumed volume of sucrose solution (*P* = 0.014) and sucrose preference in knockouts (*P* = 0.012, ANOVAs) were apparent. Such a phenotype of mild anhedonia at the beginning of sucrose-preference testing in CD2-KO mice was described previously ^24^.

### Nest building is normal in CD2-KO mice

We measured nest building as a compound indicator of hippocampus-dependent social and motivational functioning ^30^. Neither the nest score nor the amount of untorn material (bits of >0.1 g) were significantly different between genotypes (*P* > 0.2 for within-gender comparisons of both parameters, Mann-Whitney-U-test, Figure 5h-j).

### CD2-KO mice show enhanced reciprocal social interaction

Test mice were left to interact with younger, same-sex stimulus mice in novel cages. Surprisingly, the total interaction time and the number of interactions were significantly higher in CD2-KO mice compared to wildtypes (*P*_genotype_ < 0.05, ANOVA; Figure 5k-l). Males showed more interaction than females (P_sex_ < 0.0005 for both parameters), and the increased interaction in CD2-KO mice reached significance only in the male subgroup in both parameters (*P*_males_ < 0.02, *P*_females_ > 0.1, *t*-tests). This difference was mainly driven by the early stages of the test, in which CD2-KO mice showed a sustained high level of interaction time (*P*_interval*genotype_ = 0.003, repeated-measures ANOVA; Figure 5n).

## DISCUSSION

We here did a comprehensive behavioural assessment of deficits across the three symptom domains of schizophrenia in the Cyclin-D2-KO mouse model of ventral hippocampal hyperactivity. It was shown previously, that this model gains its construct validity from (1) a hypofunction of Parvalbumin-positive interneurons, (2) reduced hippocampal volume, and (3) a lack of adult neurogenesis, three cellular pathologies observed in schizophrenia ^2,18,19,23–25,37^. It also displays a mesolimbic hyperdopaminergic state, behavioural hyperactivity, and elevated activity of hippocampal subfields CA1–3, all of which could be ameliorated by increasing inhibition in the ventral hippocampus ^18^. This previous work suggests that CD2- KO-mice could provide a unique rodent model to understand the relationship between the endophenotype of hippocampal hyperactivity - analogously observed in patients with schizophrenia at baseline ^2,3,38–41^ and during sensory processing ^42,43^ - and symptoms of this disease. Hippocampal hyperactivity correlates with positive, but also with cognitive symptoms in patients, while conflicting results exist regarding the negative symptom domain ^3,41^.

Regarding cognitive symptoms, our data demonstrates impairments of CD2-knockouts in cognitive flexibility, affecting both rule-shift and rule-reversal learning (extending earlier findings, see Table 1), and uncovers a previously unknown impairment in spatial working memory. The latter aligns with a negative correlation between hippocampal hyperactivity and working memory seen in schizophrenia patients ^41^, and with impairments of short-term memory produced by broad chemogenetic ^15^ or pharmacological ^16^ hippocampal disinhibition in rodents. In contrast to patient data (and the just-cited rat study), however, we could not find an indication that CD2-KO mice are impaired in sustained attention or short-term habituation of stimulus-specific attention. Notably, the latter has been suggested to underlie aberrant salience ^33^, and thereby psychosis ^44^, which implies that these mice probably do not model the psychotic state of schizophrenia. This aligns with their profile of ventral hippocampal hyperactivity (see Introduction) and with their normal response to amphetamine (however, see ^18^).

Nevertheless, CD2-KO mice did display robust novelty-induced hyperlocomotion, which is usually considered a rodent correlate of the positive symptom domain (note however, that individual positive symptoms can appear in prodromal patients ^45^, and that the extent of hyperlocomotion in this mouse line appears considerably smaller than in a mouse model with impaired short-term habituation, the *Gria1*-knockout mouse ^33,34,46^). Interestingly, the relative elevation of locomotion compared to controls was maintained even after treatment with D1- or D2-receptor antagonists, in line with the unresponsiveness of baseline hippocampal hyperactivity to antipsychotic medication in patients ^3^. In contrast, novelty-induced hyperlocomotion was completely normalized by the mGlu2/3-agonist LY379268, supporting the notion, that hippocampal glutamatergic activity can be scaled down by this pharmacological intervention ^2^.

Furthermore, CD2-knockouts did not display robust deficits in the negative symptom domain, including social interaction, nest building, anhedonia, or incentive motivation, nor altered motor impulsivity, anxiety or perseveration. Generally, their deficit profile resembles schizophrenia - especially its prodrome - more than any other psychiatric disorder.

Based on the presented phenotyping, this mouse model appears well-suited to develop therapeutical interventions targeted at cognitive deficits caused by an overactive hippocampus, and, more generally, at halting the transition from the prodromal to the psychotic state ^4^.

## Acknowledgments

***Funding:*** This work was funded by the Institute of Applied Physiology (Ulm University), and by grants from the Juniorprofessorship programme of Baden-Württemberg (D.K.) and the Else-Kröner- Fresenius/German-Scholars-Organization Programme for excellent medical scientists from abroad (D.K.). ***Author contributions:*** C.M.G., S.A., St.S. and D.K. did experiments. J.T. programmed operant box paradigms. P.S. and B.L. contributed essential resources. D.K. designed the study and wrote the manuscript, which was revised by all authors. ***Acknowledgments:*** We thank Bastiaan van der Veen and Kasyoka Kilonzo for logistical support, Chris Barkus for providing Med-PC code, and Alexa Hagedorn, Ekaterina Merkel and Olena Sakk for assistance with colony maintenance, the animal licence and related animal work.

